# Molecular Dynamics Simulations Show How Antibodies May Rescue HIV-1 Mutants Incapable of Infecting Host Cells

**DOI:** 10.1101/2022.09.02.506285

**Authors:** Dharanish Rajendra, Nikhil Maroli, Narendra M Dixit, Prabal K Maiti

## Abstract

High mutation and replication rates of HIV-1 result in the continuous generation of variants, allowing it to adapt to changing host environments. Mutations often have deleterious effects, but variants carrying them are rapidly purged. Surprisingly, a particular variant incapable of entering host cells was found to be rescued by host antibodies targeting HIV-1. Understanding the molecular mechanism of this rescue is important to develop and improve antibody-based therapies. To unravel the underlying mechanisms, we performed fully atomistic molecular dynamics simulations of the HIV-1 gp41 trimer responsible for viral entry into host cells, its entry-deficient variant, and its complex with the rescuing antibody. We find that the Q563R mutation, which the entry-deficient variant carries, prevents the native conformation of the gp41 6-helix bundle required for entry and stabilizes an alternative conformation instead. This is the consequence of substantial changes in the secondary structure and interactions between the domains of gp41. Binding of the antibody F240 to gp41 reverses these changes and re-establishes the native conformation, resulting in rescue. To test the generality of this mechanism, we performed simulations with the entry-deficient L565A variant and antibody 3D6. We find that 3D6 binding was able to reverse structural and interaction changes introduced by the mutation and restore the native gp41 conformation. Viral variants may not only escape antibodies but be aided by them in their survival, potentially compromising antibody-based therapies, including vaccination and passive immunization. Our simulation framework could serve as a tool to assess the likelihood of such resistance against specific antibodies.

## 1 Introduction

The human immunodeficiency virus type 1 (HIV-1) has high *in vivo* mutation and replication rates, resulting in the continuous generation of mutant strains that enable escape from host immune responses and drugs.^1, 2^ Combination antiretroviral therapy limits the emergence of such resistance by increasing the genetic barrier to resistance. ^3, 4^ Antibody-based treatments have shown promise with reduction of viral loads soon after administration.^5–7^ Resistance to antibodies, however, leads to an eventual rebound of viral loads and can compromise vaccines and passive immunization strategies aimed at achieving long-term HIV-1 remission or prevention of transmission.^8–10^ Efforts are underway to devise combination therapies involving antibodies that may overcome such resistance. ^11^ Studies are also investigating the effectiveness of certain dendrimers in preventing sexual transmission of HIV-1.^12, 13^

Resistance mutations often incur fitness costs. ^14, 15^ Mutations with significant deleterious effects are naturally purged and thus are thought of as not capable of driving resistance. In an interesting recent observation, an HIV-1 variant that could not infect cells was found to have its infectivity restored in the presence of specific antibodies that target HIV-1.^16^ This implied that antibodies could effectively “increase” the fitness of mutants. Such mutants could become a source of sustained viral replication even in the presence of antibodies and create the opportunity for the development of resistance. It is important, therefore, to understand the mechanism(s) of such antibody-driven restoration of infectivity and assess whether it occurs more widely. Here, we employed fully atomistic molecular dynamics (MD) simulations to address this question.

HIV-1 entry into host cells is driven by its envelope (Env) spike protein, a trimer of non-covalently attached extracellular gp120 and transmembrane gp41 subunits. Gp120 binds to the CD4 receptor on CD4^+^ T-cells,^17^ following which Env undergoes a series of conformational changes enabling binding of gp120 to the co-receptor CCR5 or CXCR4 and its detachment from the gp41 stem.^18^ This exposes the fusion peptide (FP) of gp41, which penetrates and anchors itself into the T cell membrane. ^19, 20^ The linear pre-fusion (PrF) complex of the gp41 trimer then undergoes a large conformational change involving hairpin bends at the immunodominant loops (IL) and transitions into the 6-helix bundle post-fusion (PoF) complex (Fig. 1). This “folding-in” of the complex brings the viral and T cell membranes into close proximity and facilitates membrane fusion, and viral entry into the host cell. ^21^ Thus, stability of the PoF complex is essential for gp41 to be able to hold the membranes in proximity for the few microseconds required for the onset of fusion.^21^ Several computational studies employing molecular dynamics simulations have also contributed to our understanding of the process.^22–24^ Computational investigations have also illuminated the molecular mechanism of action of dendrimers in preventing viral transmission aiding in their further development.^25, 26^

**Figure 1:**
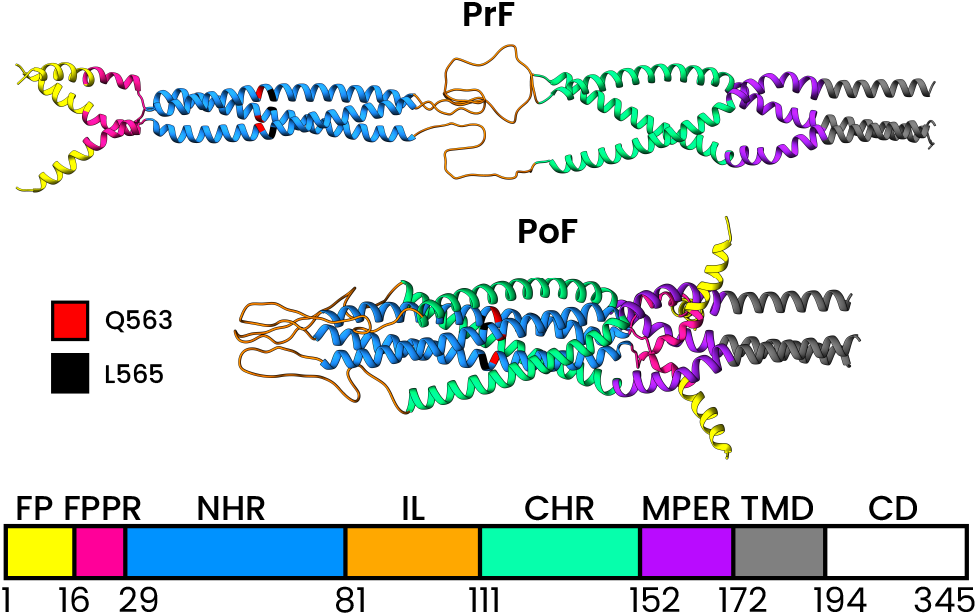
Structure and sequence of gp41 and the residues mutated. Structure of the pre-fusion (top) and post-fusion (middle) conformations of the gp41 trimer. The individual domains on the gp41 monomers are colored according to the regions demarcated by the residue numbers in the gp41 sequence (bottom): Fusion Peptide (FP), Fusion Peptide Proximal Region (FPPR), N-terminal Heptad Repeat (or Helical Region) (NHR), Immunodominant Loop (IL), C-terminal Heptad Repeat (or Helical Region) (CHR), Membrane Proximal External Region (MPER), Transmembrane Domain (TMD), Cytoplasmic Domain (CD). The residues Q563 and L565 (numbered according to the gp160 sequence) in the NHR, where the mutations are performed, are highlighted in different colors.

Joshi et al.^16^ found that the variant with the loss of infectivity harbored the Q563R mutation, which agrees with the deleterious nature of Q563R found previously. ^27^ Their study argued that the mutation affected the formation of the 6-helix bundle and compromised infectivity. Antibodies that bind cluster-I epitopes on gp41 (cluster-I antibodies) restored infectivity. These cluster-I Abs are non-neutralizing,^28, 29^ but can recognize HIV-1 infected cells and trigger antibody-dependent cell-mediated cytotoxicity.^30^ These antibodies are also involved in the inhibition of HIV replication in macrophages and immature dendritic cells. ^31^ The F240 antibody (Ab) in particular has gained a lot of attention are there have been many ideas and attempts at using it to develop a cure or a prophylactic.^32–34^

While there exist a few computational studies to uncover the mechanisms of resistance of various mutations against fusion inhibitors,^35–37^ a molecular understanding of these observations of loss of infectivity and rescue, however, is lacking. Here, using known molecular models of the gp41 trimer and the cluster-I Ab F240, we performed fully atomistic MD simulations to unravel the molecular and structural underpinnings of antibody-dependent rescue of infectivity. We show that the Q563R mutant gp41 trimer adopts not the native 6-helix bundle conformation but an alternative “bent-over” conformation due to changes in the secondary structure as well as interactions between the domains of gp41. The binding of F240 Ab to the gp41 trimer reverses these changes and restores the native conformation, thus resulting in the rescue of infectivity. Next, to test the generality of the principle of restoration of the gp41 structure with deleterious mutations by host antibodies, we repeated the simulations with another mutation and antibody pair: the L565A mutant, which is entry-defective due to the loss of fusion capability of gp41,^38–42^ and the 3D6 Ab.^43^ We find similar behavior with this variant-antibody pair as well. The L565A mutation destabilizes the native 6-helix bundle conformation and stabilizes an alternative conformation; the binding of 3D6 Ab re-establishes the native conformation.

The rest of the paper is organized as follows. In Section 2 we describe the methods that were used in obtaining the protein complexes, the simulations, and the analysis of the simulation trajectories. The results in Section 3 are divided into 2 parts. Section 3.1 deals with the destabilization of the six-helix bundle form of gp41 due to mutations. Section 3.2 describes re-stabilization due to the binding of antibodies. In Section 4 we discuss the interpretations and implications of these results. We conclude in Section 5 with a summary.

## 2 Methods

### 2.1 Protein Structures

The structure of gp41 PoF (referred to as WT gp41) was taken from our earlier studies which was obtained by homology modeling using templates of SIV. ^44^ The structures of Q563R and L565A mutant were obtained by performing three mutations (Q52R, L54A respectively in each of the gp41 monomers of the six-helix bundle) to the WT gp41 structure using FoldX. ^45^ For the F240 Ab, we used crystal structure from the RCSB Protein Data Bank (accession number 5DRZ),^30^ which has a resolution of 2.54 Å. The structure of the 3D6 Ab was also obtained from RCSB Protein Data Bank (accession number 1DFB),^46^ which has a resolution of 2.70 Å.

The complexes of Q563R gp41 bound to F240 Ab and L565A gp41 bound to 3D6 Ab were obtained through docking. The docking was performed with mutant gp41 and antibody structures using ZDock server. ^47^ The epitope of F240, 3D6 Abs on gp41, as determined experimentally,^48^ and binding site on F240^30^ was used to filter out the plausible binding modes of the Q563R gp41-F240 and L565A gp41-3D6 complexes i.e. only those binding modes were retained which had interactions between the experimentally known binding sites. Following this, the three distinct binding modes with the highest docking scores were chosen for each of the complexes and simulated in water for 30 ns to observe the evolution of the binding conformation. For each complex, the one in which the binding site and the relative orientations of the two proteins remained constant over time were chosen for the final 400 ns simulation. See Movie S2 for the structural evolution of the Q563R gp41-F240 and L565A gp41-3D6 complexes.

### 2.2 Simulation Details

All the MD simulations were performed using GROMACS 2021,^49–52^ with Amber-99SBildn force fields.^53^ The proteins were placed in a periodic rhombic dodecahedron box in which each of the faces of the box was at least 1 nm away from the protein. The docked structures i.e. Q563R gp41-F240 Ab and L565A gp41-3D6 Ab complexes were subject to an initial energy minimization in vacuum for 500 steps using a steepest descent algorithm with a force tolerance of 1000 kJ mol^−1^ nm^−1^ steps to remove any close contacts at the interface of the two proteins. All proteins and vacuum energy minimized complexes were placed in the center of a truncated dodecahedral box of TIP3P water,^54^ maintaining a minimum distance of 1 nm from all sides. Na+ and Cl-ions were added in appropriate numbers to neutralize the charge of the system. The solvated systems were also energy minimized for 3000 steps using a steepest descent algorithm. The energy minimized system was heated from 0 K to 310 K within 100 ps in an NVT ensemble with position restraints on all heavy atoms of the proteins using a force constant of 1000 kJ mol^−1^ nm^−2^. A modified Berendsen thermostat (called velocity rescaling)^55^ was used for temperature regulation, with a coupling constant of 0.1 ps. After this initial NVT simulation, 100 ps long NPT simulation was done at 1 bar to get the equilibrium bulk density. During NPT simulation, all the heavy atoms of the protein were position restrained using a force constant of 1000 kJ mol^−1^ nm^−2^ as before. Parrinello-Rahman barostat^56^ with a coupling constant of 2 ps and isothermal compressibility of 4.5*×*10^−5^ was used during the NPT runs. Following this, unrestrained production MD runs for 400 ns were performed in an NPT ensemble. Trajectories were saved every 100 ps for data analysis. Particle Mesh Ewald (PME) method was used to compute the long-range electrostatic interaction with a real space cutoff of 1.0 nm and a fourier grid spacing of 0.16 nm.^57^ Leapfrog integrator with an integration timestep of 2 fs, and the LINCS algorithm^58^ to constrain bonds involving H atoms were employed during the MD simulations.

### 2.3 Trajectory Analyses

Root mean square deviation (RMSD), Radius of gyration (R*_g_*), principal component (PC) analysis, and free energy surface (FES) calculations were performed using the tools available in the GROMACS package. PCA calculates the mass-weighted covariance matrix of the trajectory of the backbone atoms for the last 200 ns of the simulation. Eigenvalues of this matrix are obtained and sorted in decreasing order of the corresponding eigenvalues. This was done using the command gmx covar. The first two eigenvectors can explain a large portion of the variation in the structure of the protein.

Then, the projection of the trajectory on these two largest eigenvectors were taken as the coordinates for FES analysis. The command gmx sham was used to generate the Gibbs free energy surfaces, which is done by generating a 2-dimensional histogram of the projections on the largest eigenvectors followed by Boltzmann inversion.

All structures corresponding to the minimum of the well in the FES of WT gp41 were averaged to obtain the average WT stable structure. This structure was then used as the reference structure for the RMSD calculation from all trajectories. The command gmx rms was used to perform the RMSD calculations after least-squares fitting to the reference structure.

End-state binding free energy calculations using MMGBSA methods were done using the tool gmx MMPBSA^59^ over the last 200 ns of the simulation, with the Amber-99SBildn force fields, a modified Generalized-Born model of implicit solvation,^60^ the interaction-entropy approximation^61^ to calculate the entropic contribution to the free energy. An internal dielectric constant of 1, external dielectric constant of 80, and surface tension of 0.0072 kcal mol^−1^ Å^−2^ were used.

Generation of protein snapshots and trajectory videos were done in UCSF Chimera.^62^ All the computed structural and thermodynamics quantities mentioned in the text are averages over the last 200 ns of the simulation (2000 frames/snapshots).

## 3 Results

The initial simulation set-up we employed is depicted in Fig. 2. The simulations are intensive and involve up to 600,000 atoms (see Methods).

**Figure 2:**
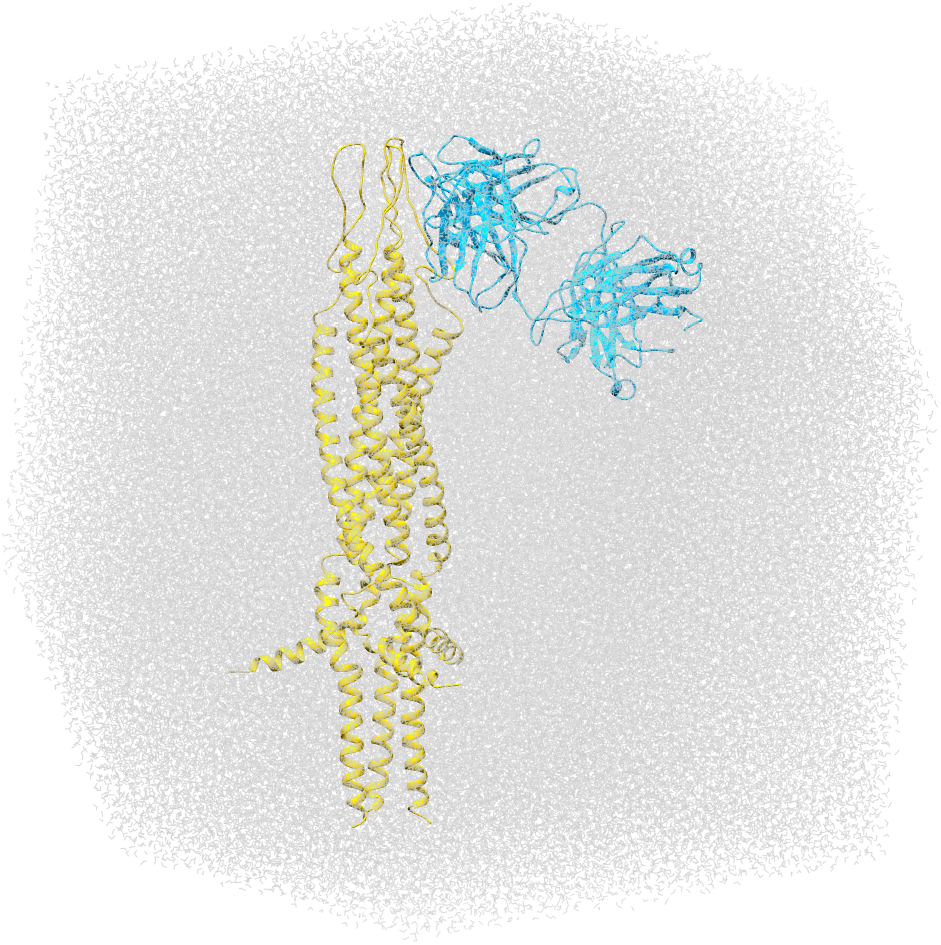
The full simulation box of the starting frame of the simulation of the complex of L565A gp41 with 3D6 antibody. Gp41 is shown in yellow, 3D6 is shown in blue and water molecules in gray. This particular truncated dodecahedral simulation box contained a total *∼*450,000 atoms. The largest system we simulated in this work consisted of *∼*600,000 atoms.

### 3.1 The Mutations Destabilize Six-Helix Bundle Form of gp41

#### 3.1.1 Structure and Shape of gp41

The final structures of the wild-type (WT) gp41 trimer and gp41 trimer with Q563R mutations (henceforth referred to as Q563R gp41) differ in one major aspect. The immunodominant loop (IL) regions are folded and lie along the axis of the helix-bundle in WT. In Q563R gp41, they are folded at a large angle to the axis of the helix-bundle in the mutant (Fig. 3B, Movie S1). This is also reflected in the radius of gyration of gp41 as shown in Fig. 4A. The radius of gyration is consistently lower for Q563R gp41 than WT. It is lower for Q563R gp41 by 7% at the maximum and by 2.6% on average gp41 (see Table 1 for the individual average values over the last 200 ns). Similarly, the L565A gp41 also adopts a conformation different from WT gp41, as seen in 3E. The IL region in L565A gp41 is slant and at a larger angle from the axis of the helix bundle. This difference in conformation is also confirmed by the consistently higher *R_g_* of L565A gp41 compared to WT gp41 (Fig. 4A).

**Figure 3:**
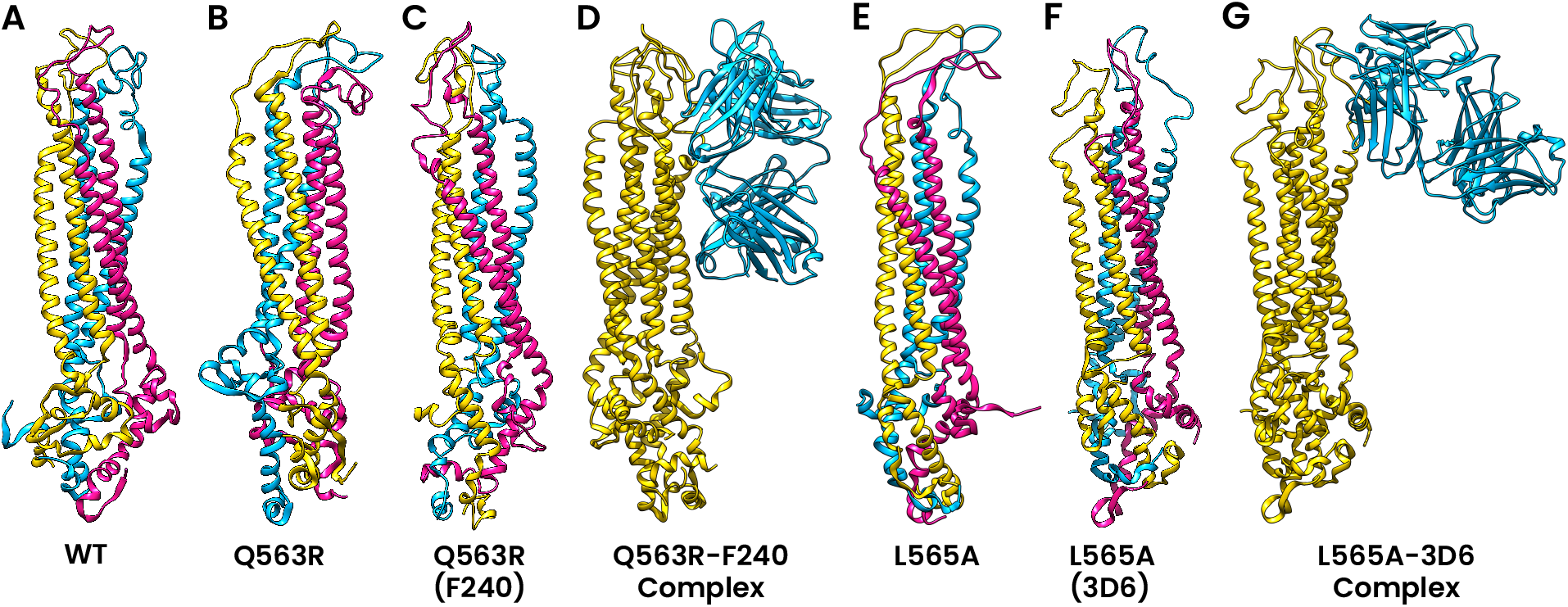
Instantaneous snapshots of the protein structures at the end of the simulations (400 ns) of (A) WT gp41 (B) unbound Q563R gp41 (C) Q563R gp41 when bound to F240 (D) complex of Q563R gp41 with F240 Ab (E) L565A gp41 (F) L565A gp41 when bound to 3D6 Ab, and (G) complex of L565A gp41 with 3D6 Ab. Unbound Q563R and unbound L565A show conformations different from the WT in which the upper immunodominant loop (IL) region bends over to one side and, is slant and at a large angle from the axis of the helix bundle, respectively. When bound to antibodies F240 and 3D6, respectively, the structures of Q563R and L565A gp41 are restored close to that of WT gp41.

**Figure 4:**
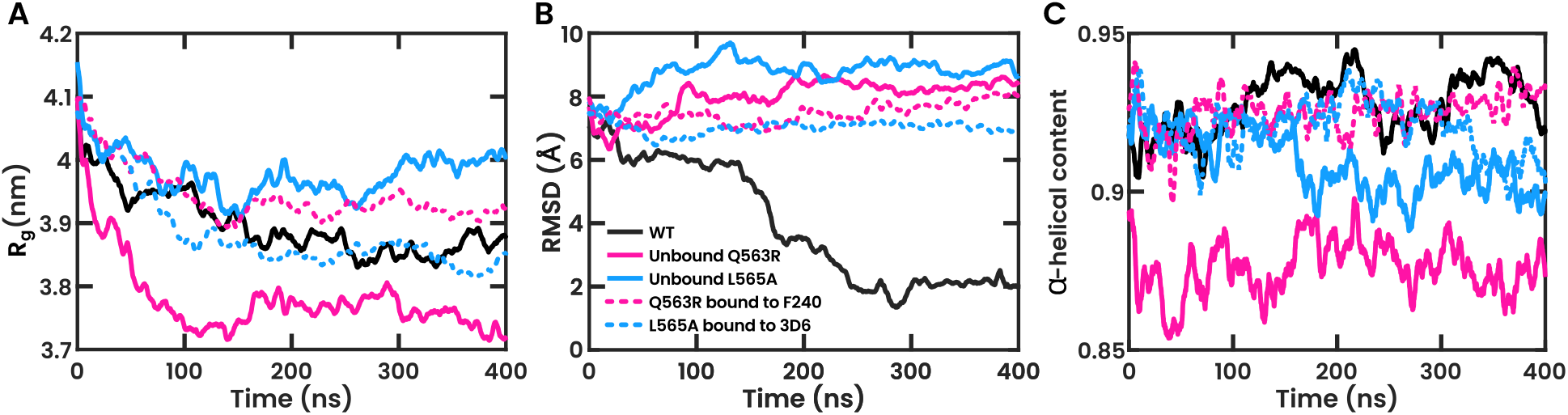
Various metrics indicating structural and conformational changes in different protein structures over the course of the 400 ns simulations.(A) Radius of gyration of gp41, (B) RMSD (root mean square deviation) of gp41 ectodomain against a reference structure of the average stable structure of WT gp41 (as determined by the FES), and (C) the fraction of residues in NHR and CHR which adopt an *α*-helical structure. The legend in (B) also applies to (A,C). The gp41 mutants deviate from WT in all these metrics. When bound to the corresponding antibodies, these metrics are restored close to that of WT. See Table 1 for a comparison of the average values of these quantities over the last 200 ns of the simulation.

**Table 1:**
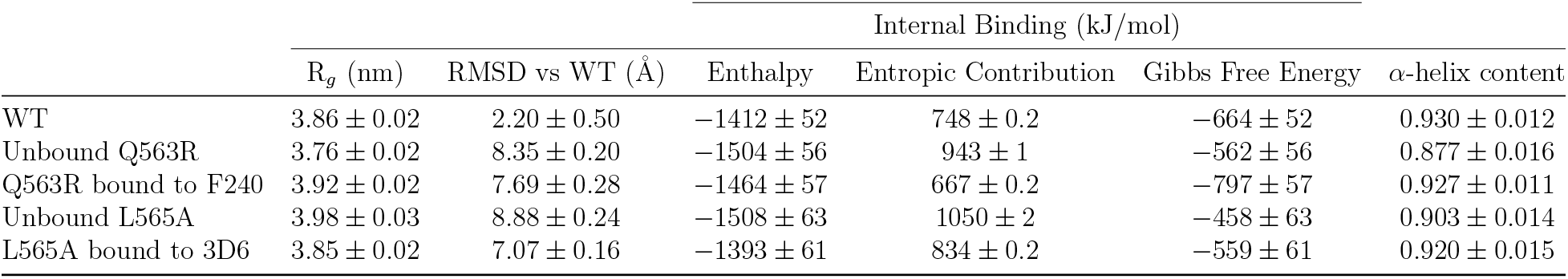
Structural and energy metrics of gp41 in its WT, mutated, and antibody-bound states estimated from simulations.

| | $R_g$ (nm) | RMSD vs WT (Å) | Internal Binding (kJ/mol) | | | |
| --- | --- | --- | --- | --- | --- | --- |
| | | | Enthalpy | Entropic Contribution | Gibbs Free Energy | α-helix content |
| WT | $3.86 \pm 0.02$ | $2.20 \pm 0.50$ | $-1412 \pm 52$ | $748 \pm 0.2$ | $-664 \pm 52$ | $0.930 \pm 0.012$ |
| Unbound Q563R | $3.76 \pm 0.02$ | $8.35 \pm 0.20$ | $-1504 \pm 56$ | $943 \pm 1$ | $-562 \pm 56$ | $0.877 \pm 0.016$ |
| Q563R bound to F240 | $3.92 \pm 0.02$ | $7.69 \pm 0.28$ | $-1464 \pm 57$ | $667 \pm 0.2$ | $-797 \pm 57$ | $0.927 \pm 0.011$ |
| Unbound L565A | $3.98 \pm 0.03$ | $8.88 \pm 0.24$ | $-1508 \pm 63$ | $1050 \pm 2$ | $-458 \pm 63$ | $0.903 \pm 0.014$ |
| L565A bound to 3D6 | $3.85 \pm 0.02$ | $7.07 \pm 0.16$ | $-1393 \pm 61$ | $834 \pm 0.2$ | $-559 \pm 61$ | $0.920 \pm 0.015$ |

#### 3.1.2 Free Energy Surface of gp41

To understand the thermodynamic origin of the difference between the WT gp41 and the Q563R and L565A mutant gp41 structures, we computed the free energy surface (FES) along the two reaction coordinates: the first and second largest principal components of gp41, using the last 200 ns of the simulations. Considerable information can be gained from an FES of this form. Neighboring points correspond to similar structures, and the structure corresponding to the minimum of a well is similar to the average structure represented by the entire well. The number of wells gives us an idea of the number of distinct conformations that are sampled by the protein. And, the size and depth of the well tells us about the relative thermodynamic stability of the different conformers.

The FES of WT gp41 contains a single deep well corresponding to its most stable conformation (Fig. 5A). In this conformation, the IL lies along the axis of the helix bundle. In contrast, the FES of Q563R gp41 shows three relatively deep wells (Fig. 5B). All three wells correspond to the bent-over conformation in which the IL region is bent at a large angle from the axis of the helix bundle, with some small variations. The native WT conformation is completely absent in the FES of Q563R gp41, indicating that it is not a stable conformation for Q563R gp41. For the case of unbound L565A gp41, the FES shows two deep wells (Fig. 5D), both quite similar and corresponding to the slant conformation in which the IL region is extended but slant at an angle from the axis of the helix bundle. The native WT gp41 conformation is absent from this FES indicating that it is an unstable conformation for L565A gp41.

**Figure 5:**
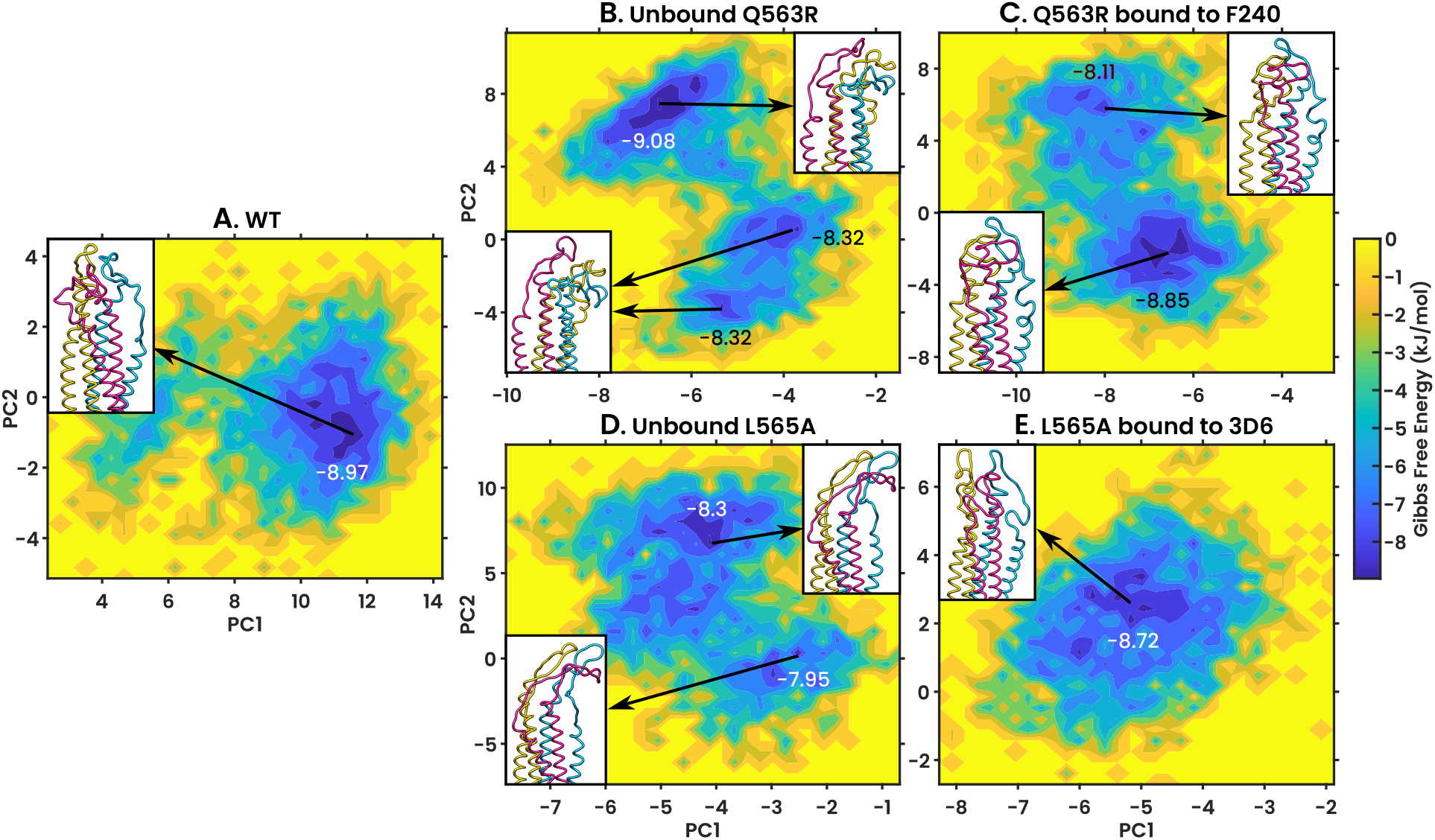
Free energy surfaces of the projections of the trajectory along the first and second largest principal components of (A) WT gp41 (B) Q563R gp41 (C) Q563R gp41 when bound to F240 Ab (D) L565A gp41 (E) L565A gp41 when bound to 3D6 Ab. The FES is calculated using configurations from the last 200 ns of the 400 ns trajectory. Insets: average (PC1 + PC2) backbone structures corresponding to the indicated minima. The maximum depth of each important well is indicated next to it in kJ/mol. Two wells in (B) correspond to structures that are very hard to tell apart just with the eye; these wells are depicted with a single structure for clarity. The FES of the unbound mutants contains deep wells corresponding to alternative conformations not seen in WT. On the binding of their respective antibodies, the native 6-helix bundle conformation becomes stable.

To understand how different the mutant structures are from the stable WT structure, we performed RMSD calculations of the whole trajectory of the mutants from the average stable WT structure. This average stable WT structure was obtained by averaging the structures that correspond to the minimum in the well of the FES of WT gp41 (Fig. 5A). These RMSDs are shown in Fig. 4B. As expected, the RMSD against the WT trajectory is low towards the end of the simulation, when the structure is stabilized. However, for the case of the unbound mutants, the RMSD from the WT stable structure is very high, indeed indicating that the structure assumed by the unbound mutants is quite different from the stable WT structure.

#### 3.1.3 Interactions between NHR and CHR Domains

The NHR and CHR domains interact with each other nearly along their entire lengths and this interaction maintains the 6-helix bundle form, preventing it from reverting to the linear form. We have calculated the binding free energy between the NHR and CHR domain using the MMGBSA method using the last 200 ns of the trajectory. The free energies were also decomposed into pairwise components to understand which interactions between the NHR and CHR play an important role. Although methods such as MMPBSA and MMGBSA have their advantages and disadvantages,^63, 64^ we use it here mainly because the size of our system makes it prohibitively computationally expensive to implement other methods such as Free Energy Perturbation and Replica Exchange Free Energy Perturbation. Further, this method allows for the pairwise decomposition of the free energy. Moreover, several studies have made use of such methods with implicit solvent models to successfully estimate protein–protein binding energies.^65–67^ MMGBSA has been successfully used even in the case of HIV gp41.^68^

In WT gp41, NHR and CHR interact strongly with a binding free energy of 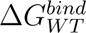 = *−*664 *±* 52 kJ/mol. The strongest interactions in WT gp41 are between

1. ARG46 on NHR and GLN139 on CHR, contributing Δ*G* = *−*54 *±* 13 kJ/mol to the total internal binding free energy, in which the oxygen atom of the side chain of GLN139 forms two H-bonds with the side chain nitrogen atom of ARG46
2. GLN66 on NHR and TRP117 on CHR, contributing Δ*G* = *−*49 *±* 7 kJ/mol
3. GLN32 on NHR and GLU342 on CHR of the next chain, contributing Δ*G* = *−*29 *±* 13 kJ/mol, with the side chain oxygen atom of GLN342 and the side chain nitrogen atom of TRP117 forming a salt bridge (according to criteria defined in^69^)
4. ARG68 on NHR and TRP306 on CHR of the next chain, contributing Δ*G* = *−*18 *±* 5 kJ/mol

The residue numbering is according to the residues on truncated gp41 trimer as shown in Fig. 1 and the numbering continues from one monomer/chain to the next.

Q563R gp41 on the other hand, has a lower binding free energy between NHR and CHR: 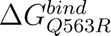 = *−*562 *±* 56 kJ/mol, 15% lower than in WT gp41. The important interactions seen in WT are weaker or nearly absent in Q563R gp41. The ARG46–GLN139 interactions contributes just Δ*G* = *−*17 *±* 14 kJ/mol, 68% lower than WT. The GLN66–TRP117 interactions contribute Δ*G* = *−*41 *±* 4 kJ/mol, 17% lower than WT. The GLN32–GLU342 interactions are almost non-existent and therefore contribute close to nothing to the total internal binding free energy. (See Fig. S1 in the supplementary information for the interaction diagrams.)

Similarly, in the L565A gp41, the binding free between the NHR and CHR domain decreased to 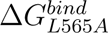 = *−*458 *±* 63 kJ/mol, 31% lower than that in WT gp41. The ARG68–TRP306 interaction is virtually absent in L565A gp41.

Therefore, the internal binding that maintains the protein structure becomes weaker due to the Q563R and L565A mutations.

#### 3.1.4 Secondary Structure of gp41

The *α* helical content of the helical regions is known to provide structural stability to the protein. The helical regions (NHR, CHR) in WT gp41 consist of 93% *α* residues. Q563R gp41 has consistently lower *α* content throughout the simulation (Fig. 4C), of about 88%. It has, at the minimum 13.2%, and on average 5.7% lower *α* content than WT gp41. (see Table 1).

Similarly, the *α*-helical content of the helical regions of L565A gp41 is consistently lower than in WT gp41 after 100 ns in the simulation, (Fig. 4C). L565A gp41 has an *α*-helix content of 90.3%. It has, at the minimum 9.5%, and on average 3% lower *α*-helical content than in WT gp41.

This loss of *α* helical content occurs at the N- and C-terminal ends of the NHR and CHR, where residues adopt bend, 3-helix, 5-helix, and unorganized coil structures (see Fig. S2 for a map of the average secondary structure across the residues of NHR, CHR). This reduction of *α* helical content and conversion to other secondary structures destabilizes not only the helical regions of Q563R and L565A gp41 but in effect the whole proteins, and results in the bending of the IL region. We next examine how the binding of the F240 and 3D6 antibodies, respectively, restores the Q563R and L565A gp41 structures.

### 3.2 Binding of the Antibodies Reverses the Changes of the Mutations

#### 3.2.1 Structure and Shape of gp41

The binding of F240 antibody (Ab) has one major effect on the structure of Q563R gp41: it restores the IL conformation to that of the WT. The IL region of Ab bound Q563R gp41 is extended along the axes of the helix bundle and not bent over like the unbound Q563R gp41 (Fig. 3C, Movie S1). The radius of gyration of Ab bound Q563R gp41 also increases and becomes close to that of WT gp41 (Fig. 4A). The radius of gyration of Ab bound Q563R gp41 is 4.3% greater than unbound Q563R gp41 and just 1.6% greater than WT gp41.

Similarly, in the presence of the 3D6 Ab, the IL region in L565A gp41 is no longer slant and now lies along the axis of the six-helix bundle, similar to WT gp41 (Fig. 3F). The restoration of radius of gyration values to those similar to that of WT gp41 (Fig. 4A) further confirm this observation.

#### 3.2.2 Free Energy Surface of gp41

In the presence of F240 Ab, the free energy landscape of Q563R gp41 changes drastically (Fig. 5C). The FES of Ab bound Q563R gp41 shows two deep wells, both corresponding to very similar structures. These structures closely resemble the native WT gp41 structure, in which the IL region is along the axis of the helix bundle. Moreover, the FES does not show the bent-over structure that is seen in unbound Q563R gp41. The RMSD of Ab bound Q563R gp41 against the WT stable structure also confirms this (Fig. 4B). This RMSD decreases for Q563R gp41 in the presence of F240 Ab by about 8%. This tells us that the structure is becoming closer to that of WT gp41, indicating that the binding of F240 Ab stabilizes Q563R gp41 in the native PoF six-helix bundle conformation. In the presence of the 3D6 Ab, the FES of L565A gp41 is modified and contains only one deep well (Fig. 5E). This well does not correspond to the slant conformation but instead to one in which the IL is along the axis of the 6-helix bundle, like in the case of WT gp41. The RMSD of Ab bound L565A gp41 against the WT stable structure also shows this. It decreases for L565A gp41 in the presence of 3D6 Ab by about 20%. This shows us that the structure becomes closer to that of WT gp41, indicating that the binding of the 3D6 Ab re-stabilizes it L565A gp41 in the native PoF six-helix bundle conformation.

#### 3.2.3 Interactions between NHR and CHR Domains

The strength of interactions between NHR and CHR domains in Q563R gp41 is increased when it is bound to F240 Ab and has a binding free energy of 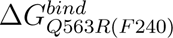 = *−*797 *±* 57 kJ/mol, 39% greater than that in unbound Q563R gp41 and is even 20% greater than in WT gp41. Here, the important interactions seen in WT gp41, which had been weakened in unbound Q563R gp41, become stronger. The interactions of ARG46–GLN139, GLN66– TRP117, and GLN32–GLU342 now have binding energies of *−*59 *±* 8*, −*58 *±* 9, and *−*25 *±* 15 kJ/mol, respectively. This brings them up to or higher than the level seen in WT gp41. These strong interactions keep the helices together and stabilize the 6-helix bundle form of gp41. (See Fig. S1 for the interaction diagrams.)

In the case of L565A gp41, when it is bound to 3D6 Ab, the interaction energy between NHR and CHR domains becomes stronger and reaches a value of 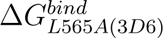 = *−*559 *±* 61 kJ/mol, 45% higher than unbound L565A. The ARG68–TRP306 interaction, which was virtually absent previously, also increases in strength to values of Δ*G* = *−*11 *±* 8 kJ/mol. This increase in the strength of binding interactions stabilizes the helix bundle and the entire protein conformation.

#### 3.2.4 Secondary Structure of gp41

The binding of the F240 Ab restores the native secondary structure in the NHR and CHR domains of Q563R and L565A gp41 (Fig. 4C). The *α* content of F240 bound Q563R gp41 is consistently higher than that of unbound Q563R gp41. It shows an increase of 5.7%, bringing it closer to that of WT gp41. The *α* content of 3D6 bound L565A gp41 also increases by 2% compared to the unbound case (Table 1). The residues at the terminal ends of the NHR and CHR are no longer in bend, 3-helix or 5-helix structures, but mostly in *α*-helix and turn structures (Fig. S2). With the restoration of the secondary structure, the 6-helix bundle regains its stability and in turn stabilizes the whole proteins. The restoration of the secondary structure also allows interactions between NHR and CHR which reinforce the stability of the proteins.

## 4 Discussion

Antibody-dependent rescue of viral variants carrying deleterious mutations ^16^ may compromise modern, antibody-based intervention strategies against HIV-1. Here, using comprehensive, large-scale MD simulations, we unraveled the molecular mechanisms underlying such rescue and argued that it is likely to be a general phenomenon, across multiple mutant-antibody pairs.

Specifically, we found that the Q563R mutation causes changes in the free-energy landscape of gp41 resulting in detrimental changes to its secondary structure in the NHR & CHR domains, and its tertiary structure of the IL region. The mutation also results in the weakening of the internal binding interactions between NHR and CHR domains. The changes in the free energy landscape indicate that the native 6-helix bundle form is no longer a stable conformation of the Q563R gp41. The free energy surface plots show that there exist other, more stable conformations that Q563R gp41 would rather adopt than the native 6-helix bundle conformation. The bent-over conformation might be among the more stable states. Further, when Q563R gp41 is bound to the F240 Ab, these detrimental changes are reversed and the free energy landscape is changed again in such a way that the native 6-helix bundle form becomes stable. F240 Ab bound Q563R gp41 resembles the WT gp41 in structure. This restorative effect of F240 Ab can be generalized to other cluster-I antibodies such as 240-D and 246-D as they have highly similar epitopes on gp41 and would induce the same effect in the protein. These observations were also reflected with a second mutant-antibody pair we studied — L565A gp41 and 3D6 Ab — arguing for the generality of the phenomenon.

The free energy surface provides further insights into the role of the rescuing antibodies. Even though the 6-helix bundle form of mutant gp41 is unstable, transitions to this conformation are still possible, following the detachment of gp120 from gp41. Due to its instability, the protein may be unable to hold the viral and T-cell membranes in proximity for the *∼*1.5 microseconds required for the fusion to begin.^21^ However, the mutant gp41 protein may still remain in the unstable 6-helix bundle conformation long enough for the antibody to bind to it. Upon binding, the structure is stabilized and maintained for an adequate duration for membrane fusion, due to the long dissociation times of antibodies.^70, 71^

It is possible that such gp41 mutants are present widely and depend on antibodies for their infectivity. Mutants that utilize and depend on broadly neutralizing antibodies (bNAbs), which are under development for treatment and prevention,^8, 10^ may also exist. Administration of bNAbs may select for these antibody-dependent resistant strains. These strains may then survive and proliferate while the bNAbs remain in circulation. In the process, they may allow resistant strains to emerge, compromising treatments. Antibody-dependent rescue of viral infectivity may thus have to be accounted for in designing antibody-based interventions and care has to be taken before using them as a widespread treatment regime.

Our simulation framework can be utilized to predict the possibility of antibody-dependent rescue for specific bNAbs. One can perform large-scale simulations of a large number of mutations of gp41 in its 6-helix bundle form. If some mutation results in destabilization of the protein, it could become a candidate for rescue. The mutant could then be examined while bound to the bNAb in question. If the protein becomes stable again, then antibody-dependent rescue and subsequent resistance may be likely. BNAbs may thus be screened and those least likely to fail and chosen for further development.

In summary, our simulations help us better understand HIV-1 and other viruses, by shedding light on the underlying mechanisms of viral proteins, and our framework may be used to aid in the eradication of the diseases caused by them.

## 5 Conclusions

In this study, we performed fully atomistic molecular dynamics simulations of the HIV-1 gp41 trimer, its entry deficient variants, and their respective complexes with antibodies. We show that the Q563R mutation effects some substantial changes in the trimer, such as a decrease in its *α*-helical content, and the weakening of the interactions between the domains of gp41. This changes the free energy landscape of the protein leading to the loss of the native WT conformation (responsible for infection) and the stabilization of an alternative conformation. Upon binding of the antibody to the Q563R gp41, these changes are reversed — *α*-helical content is restored and the inter-domain interactions are strengthened. This stabilizes and restores the native WT conformation in the mutant protein. We finally establish the generality of antibody-dependent rescue of viral variants by demonstrating the same phenomena in the case of an infection deficient L565A gp41 variant and 3D6 antibody. The L565A gp41 shows similar signs of loss of *α*-helical content and weakening inter-domain interactions. This again leads to a change in the free energy surface, loss of native WT conformation, and stabilization of an alternative conformation. A new antibody 3D6, upon binding to the infection-deficient L565A gp41, reverses these changes and restores the native WT conformation. We believe that this indicates a rescue of the L565A gp41 by 3D6 Ab. Antibody-dependent rescue of viral variants can pose a threat to antibody-based treatment and this must be taken into account in the design of these therapies.

## 6 Supplementary Information Description

The Supporting Information contains two additional figures: Fig. S1, which shows the interaction diagrams of the main interactions between NHR and CHR on WT gp41, and Fig. S2 which shows the map of the secondary structure adopted by each residue in the helical regions and its evolution over the simulation time. Additionally, there are two supporting movies: Movie S1, which shows the 400 ns trajectory of the WT gp41 protein and the mutant gp41 proteins unbound and bound to their respective antibodies, and Movie S2, which shows the 400 ns trajectory of the two mutant-antibody complexes.

## 8 Acknowledgment

DR thanks KVPY, DST India for scholarship. PKM thanks DST, India for financial support.

