## Supplementary Information for "Molecular Dynamics Simulations Show How Antibodies May Rescue HIV-1 Mutants Incapable of Infecting Host Cells"

### This PDF file includes:

Figures S1, S2
Legends for Movies S1, S2

### Other supplementary materials for this manuscript include the following:

Movies S1, S2

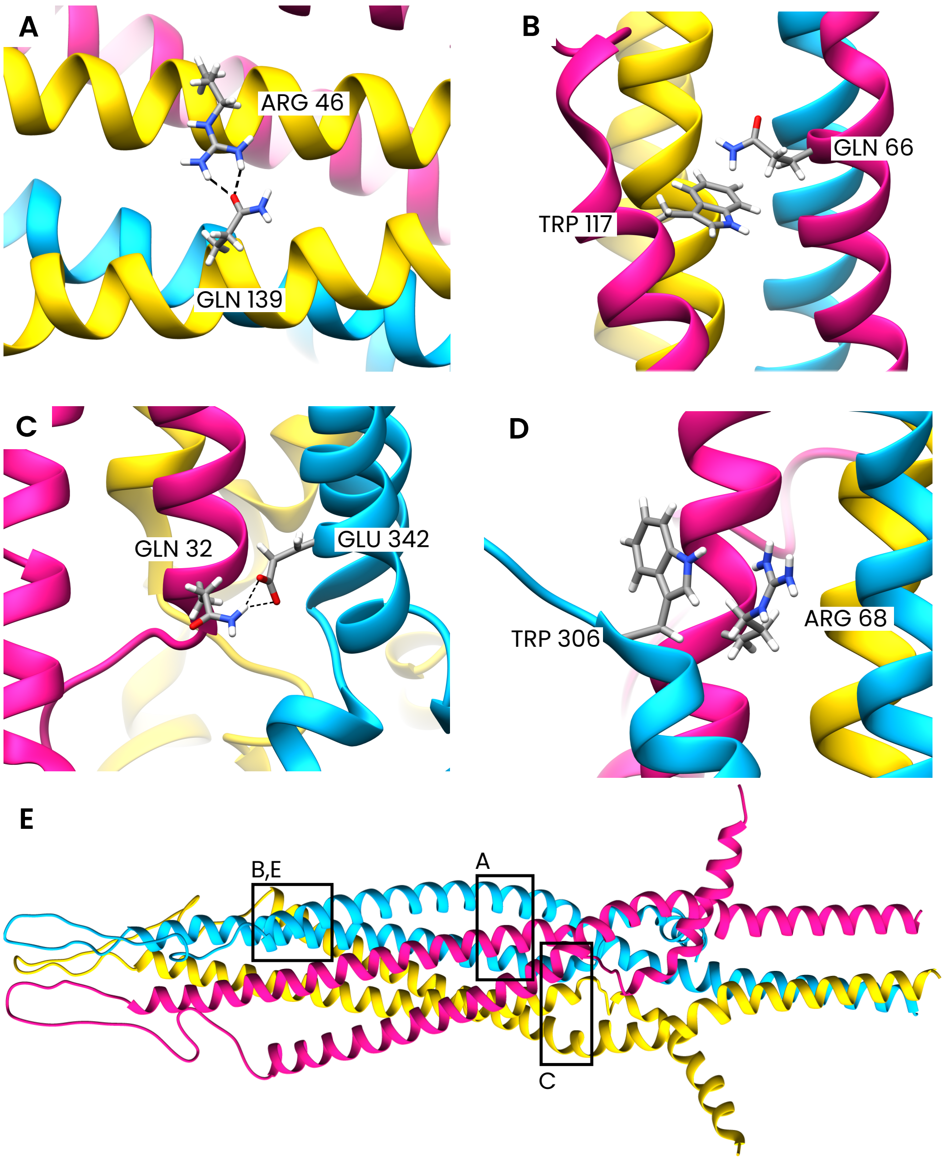


**Fig. S1.** (A-E) Interaction diagrams between residues on NHR and residues on CHR and (F) their broad locations on the gp41 trimer. Dashed lines indicate strong polar/hydrogen bonding interactions. The names of the residues are indicated next to each of the interacting residues. The three chains (monomers) of gp41 are colored differently. The side chains are colored by elements. C is in gray, H is in white, N is in blue and O is in red. All diagrams are from the last frame (400 ns) of the WT gp41 simulation. The interactions between these amino acids in the unbound and bound mutant proteins are qualitatively similar (not shown).

 

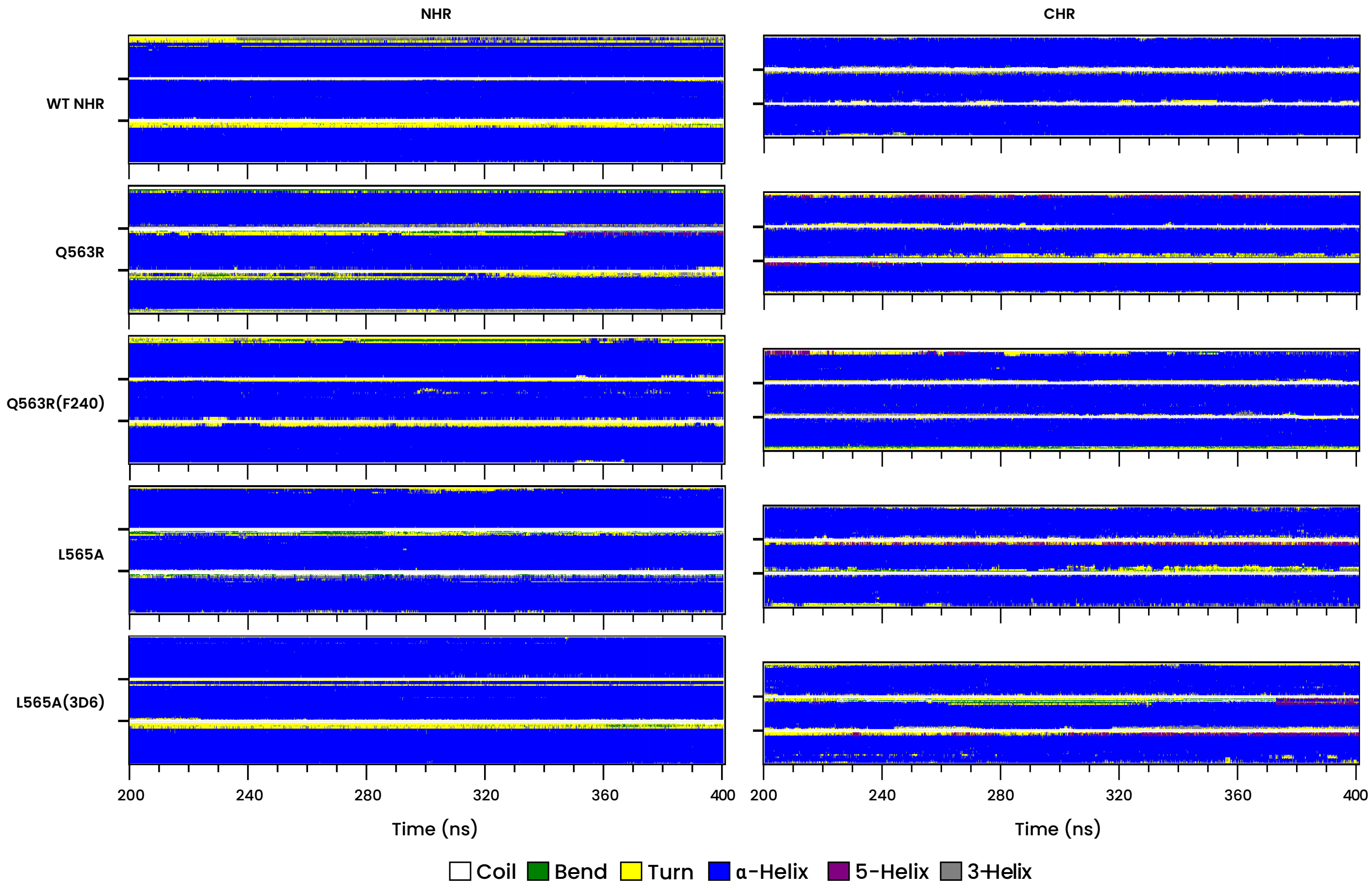


**Fig. S2.** Map of secondary structure adopted by each residue in the NHR and CHR helical regions of WT, and bound and unbound mutants over the last 200ns of the simulation. The different residues are shows across the y-axis and the two ticks on the y-axis separate the three chains of gp41. The central regions of NHR and CHR of each chain display an $\alpha$-helix structure. At their terminal ends, the residues show other structures such as bend, 5-helix and 3-helix. The width of the region of residues displaying these alternate structures increases in Q563R and is reduced by binding of F240 as shown in the map labelled Q563R(F240). However, there is not much difference in the case of unbound and bound L565A gp41.

**Movie S1. Trajectory of (A) WT gp41 (B) Unbound Q563R gp41 (B) Q563R gp41 when bound to F240 Ab (B) Unbound L565A gp41 (B) L565A gp41 when bound to 3D6 Ab projected onto their respective two largest principle components over the course of the 400ns NPT simulation. The three chains (monomers of gp41) are colored differently.**

**Movie S2. Trajectory of (A) Q563R gp41-F240 Ab complex (B) L565A gp41-3D6 Ab complex over the course of the 400ns NPT simulation. The gp41 protein are colored gold and the antibodies are colored blue.**
